# Facile detection of peptide-protein interactions using an electrophoretic crosslinking shift assay

**DOI:** 10.1101/2024.04.22.590415

**Authors:** Benjamin W. Parker, Eric L. Weiss

## Abstract

Protein-protein interactions with high specificity and low affinity are functionally important but are not comprehensively understood because they are difficult to identify. Particularly intriguing are the dynamic and specific interactions between folded protein domains and short unstructured peptides known as short linear motifs (SLiMs). Such domain-motif interactions (DMIs) are often difficult to identify and study because affinities are modest to weak. Here we describe “electrophoretic crosslinking shift assay” (ECSA), a simple in vitro approach that detects transient, low affinity interactions by covalently crosslinking a prey protein and a fluorescently labeled bait. We demonstrate this technique on the well characterized DMI between MAP kinases and unstructured D-motif peptide ligands. We show that ECSA detects sequence-specific micromolar interactions using less than a microgram of input prey protein per reaction, making it ideal for verifying candidate low-affinity DMIs of components that purify with low yield. We propose ECSA as an intermediate step in SLiM characterization that bridges the gap between high throughput techniques such as phage display and more resource-intensive biophysical and structural analysis.

## Introduction

Short linear motifs, or SLiMs (1) are unstructured peptide motifs that interact with specific binding sites on folded protein domains, frequently with low to moderate affinity (1-300 μM). Such domain-motif interactions (DMIs) are a type of dynamic protein-protein association important in cellular processes such as signal transduction and protein complex assembly through multivalent contacts that can be exploited by cellular pathogens (2). The low affinity of DMIs often makes them more difficult to detect and validate than higher affinity interactions between folded proteins, thereby potentially resulting in underrepresentation in current protein-protein interaction networks.

Straightforward pulldown or surface absorption assays are unreliable for validation of low affinity DMIs because protein-protein interactions (PPIs) are lost during wash steps due to non-equilibrium binding conditions (3). Instead, characterization of low affinity interactions between peptide ligands and folded domains generally involves relatively more laborious biophysical analyses such as fluorescence polarization (FP), isothermal calorimetry (ITC), or microscale thermophoresis (MST) detect and study these interactions, which also often require expensive reagents and specialized equipment. Thus, there is a need for rapid and technically simple approaches for identification and analysis of DMIs with micromolar affinity.

Here we present electrophoretic crosslinking shift assay (ECSA), a simple and sensitive technique for detecting weak and transient PPIs, in particular DMIs, using purified component proteins. ECSA works by detecting the in-gel shift of a GFP-tagged bait SLiM peptide ligand following in vitro chemical crosslinking to a purified folded prey domain (Figure 1A). A commonly available imidoester-based crosslinker is used to convert weak DMIs into covalent bonds. As ECSA uses two purified protein components expressed in cells, peptide synthesis is unnecessary. This technique can qualitatively detect DMIs with affinities >1 μM using less than 1 μg of purified prey protein per reaction. It can be completed in less than a day and requires only standard molecular biology reagents and commonly available gel imaging systems.

**Figure 1:**
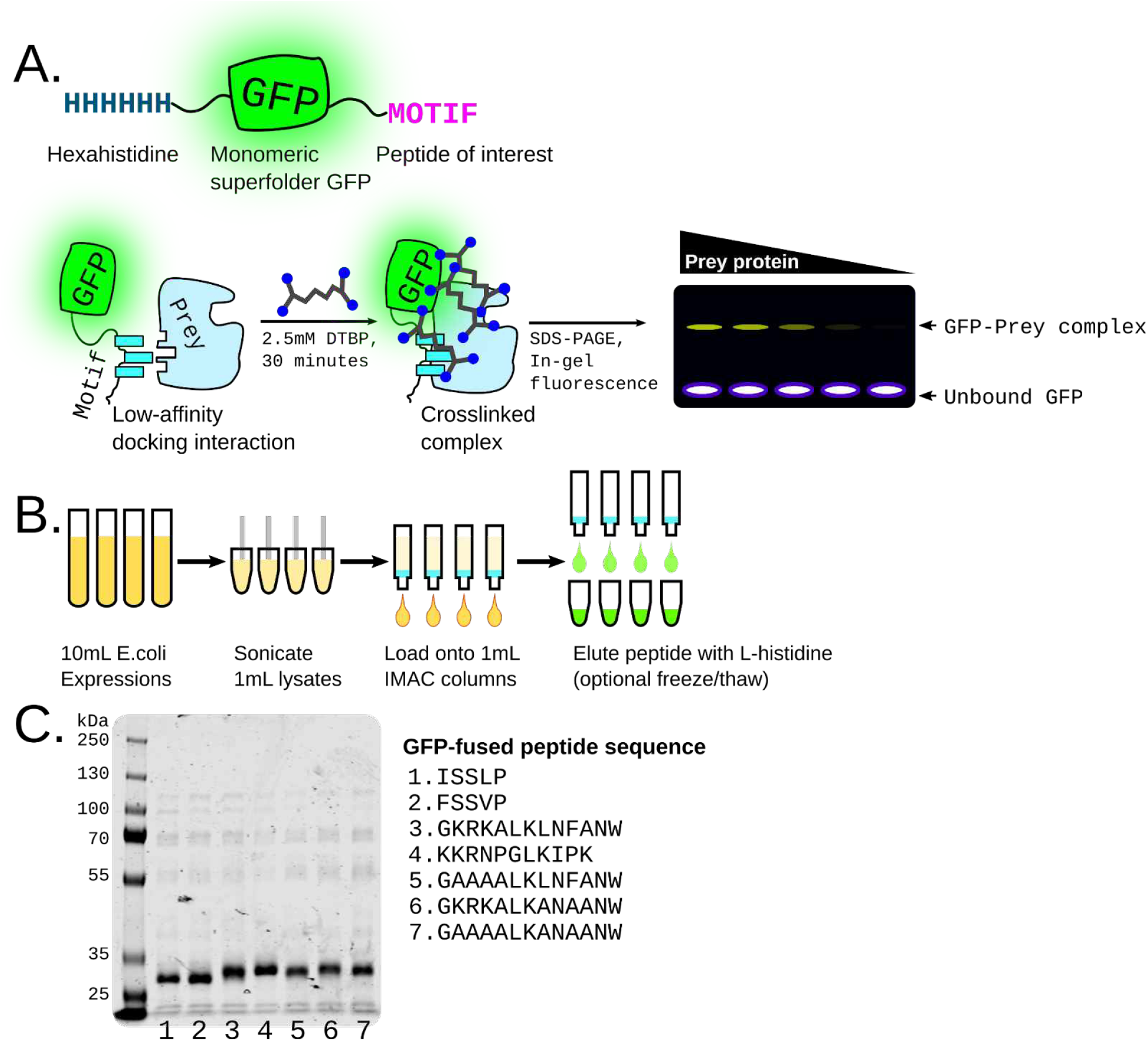
ECSA design and workflow. A, basic design of msGFP-peptide fusions used as fluorescent bait. The motif of interest is fused to monomeric superfolder GFP (9) with an N-terminal hexahistidine tag. Incubation of this motif with a prey protein in the presence of an imidoester crosslinker such as DTBP creates a crosslinked complex which can be detected by an in-gel fluorescence shift. B, workflow for purification of GFP-motif fusions in parallel. C, example of 7 peptides fused to GFP as in A purified and run on Coomassie-stained SDS-PAGE. Expected molecular weight is the range of 28-30kDa.

We demonstrate the utility of ECSA using well characterized, functionally important peptide – folded domain interactions: association of JNK1 to the MKK4 D-motif and p38a with the MKK6 D-motif, which have previously determined Kd values of ∼3.5 μM and ∼7.51 μM, respectively (4). We provide a protocol for semi-high-throughput purification of the GFP-fused bait peptide as well as for the crosslinking assay itself. We show that mutagenesis of basic and hydrophobic regions that have previously been shown to be important for D motif - kinase domain binding clearly reduces in GFP fusion protein gel shift, indicating that ECSA is a facile method for defining binding determinants in SLiM peptide ligands.

## Materials and Methods

### Expression of GFP-motif fusions

Figure 1B shows general workflow, in which peptide motifs of interest are fused C-terminally to hexahistidine-tagged monomeric superfolder green fluorescent protein (msGFP). We obtained peptide motif coding sequence as single-stranded DNA oligonucleotides (IDT, inc), which we annealed and ligated into a pBH4-based vector (5) C-terminal to msGFP. The construct was transformed into BL21 (DE3). 10mL of terrific broth was inoculated with an overnight culture of the vectors, grown at 37C for 2 hours, then expression induced with 0.2mM IPTG and growth continued at 16C overnight.

### Procedure for the purification of GFP-motif fusions

We resuspended green-hued cell pellets in 0.8mL Lysis Buffer (50mM Tris pH 8.0, 2M NaCl) containing 1mM freshly-added PMSF, lysed cells by sonication with a microtip, and spun the resulting lysate in a benchtop microcentrifuge for 15 minutes. We transferred each supernatant to a separate tube containing 40 μL of 50% Ni-NTA resin (Genscript), nutated for 20 minutes to bind GFP fusion proteins, then transferred the resin suspension to a 1 mL spin column (G-Biosciences). We collected msGFP-bound resin be spinning at 1000rcf, discarded the flow-through, washed the resin once with 0.5 mL Lysis Buffer and once with 0.5 mL Wash Buffer (100mM sodium phosphate, 50mM sodium chloride, 20mM imidazole, pH 7.4) by spinning at 1000rcf. After a brief spin of the resin alone, we eluted bound protein adding 20 μL 200mM L-histidine (pH 7.8), waiting 10 minutes, and spinning at 1000rcf for 2 minutes.

As an optional but recommended polishing step we froze the lysate at -20C and thawed at room temperature to enhance precipitation of residual contaminating proteins, then spun down at max speed. Resulting purified msGFP-peptide fusion can be frozen at -20C in the 200mM L-histidine elution buffer for >3 years.

### Procedure for the purification of JNK1 MAP kinase

We purified hexahistidine-tagged JNK1 as previously described (5).

### Procedure for the crosslinking assay

We quantified msGFP-peptide fusion concentration by A280 and adjusted to 50 μM (50pmol/ μL), diluting into 200mM L-histidine. Hexahistidine-tagged JNK1 was exchanged into kinase buffer (20mM HEPES pH 8.0, 150mM sodium chloride) using Sephadex G-25 gel filtration resin (Cytiva). Bovine serum albumin (BSA) (Sigma) was dissolved in kinase buffer to 20mg/mL. To crosslink, we added 0.5 μL of 50 μM msGFP-peptide fusion to 3.5 μL of JNK1 or BSA. We then dissolved the crosslinker dimethyl 3,3’-dithiobispropionimidate (DTBP; Thermo Scientific Pierce) in kinase buffer to final concentration of 25mM and immediately added and 0.5 μL to the protein solution, to a final volume of 4.5 μL with final concentrations of DTBP and msGFP-peptide fusion at 2.8mM and 5.6 μM, respectively. We allowed crosslinking to proceed at room temperature for 35 minutes, and then stopped the crosslinking reaction by adding 10 μL of quenching/loading buffer (1M glycine, 20% glycerol) with gentle mixing. We loaded reactions onto a 12% tris-glycine SDS-PAGE gel and separated at 200V for 35 minutes and imaged gels were imaged using a Sapphire image (Azure Biosystems) using the SmartScan option.

### Gel shift image processing

We processed TIFF format raw images obtained from the Sapphire platform with the ImageJ gel analysis toolset (6). To make figures 2 and 3 we used the ImageJ “fire” look up table to more clearly visualize signal intensity; to produce high contrast images for figures (denoted †) we contrast enhanced the raw image was using intensity histogram equalization. We used ImageJ’s gel analysis package on the original (unmodified) TIFF image to quantify binding.

**Figure 2:**
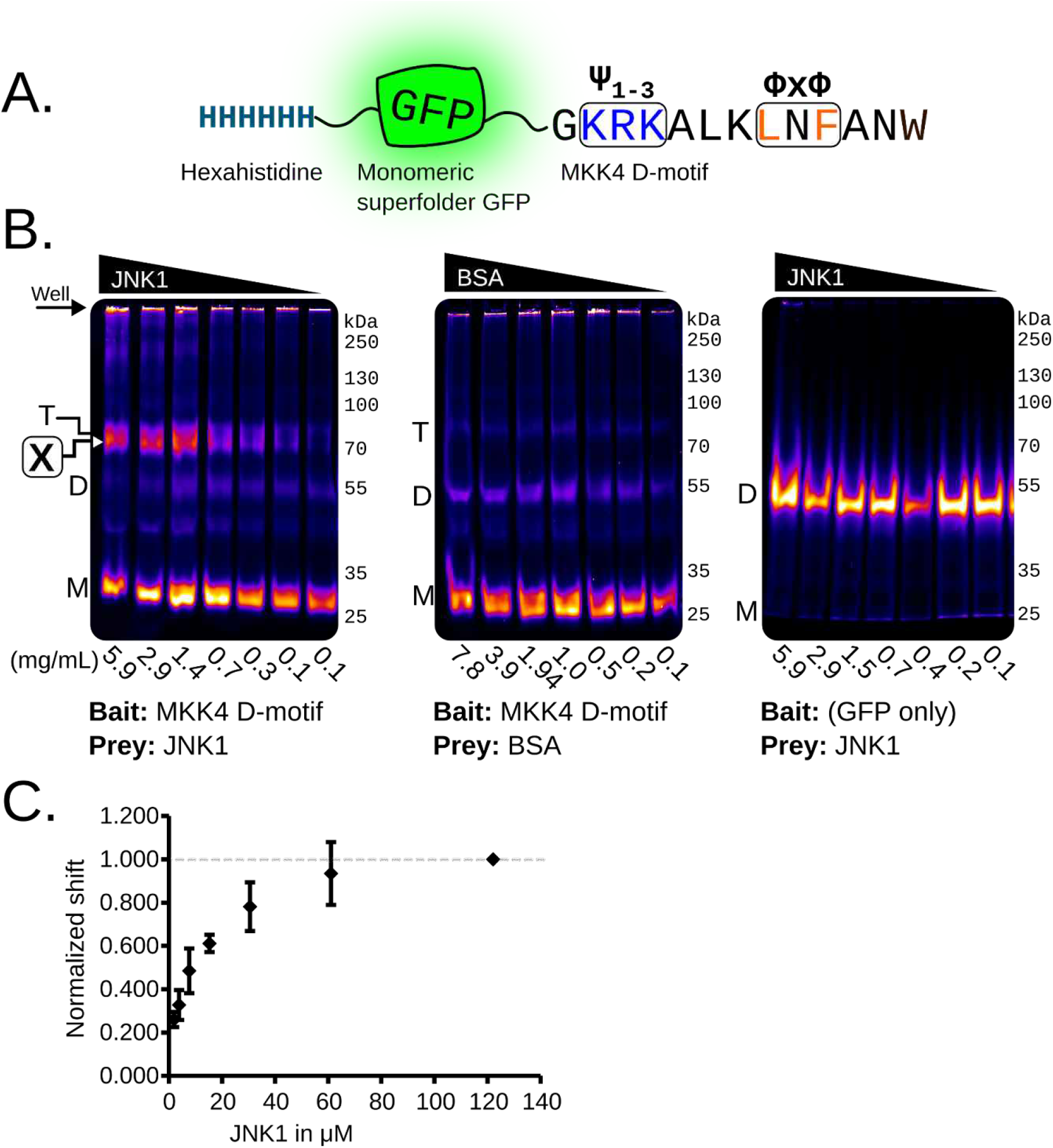
ECSA confirms the well-characterized interaction between the MKK4 D-motif SLiM and the MAP kinase JNK1. A, msGFP-MKK4 construct used as the bait peptide. The canonical basic (Ψ1-3) and hydrophobic (ΦxΦ) regions are boxed and highlighted as blue and orange, respectively. B, ECSA gel shift demonstrating binding curve with decreasing concentrations of purified JNK1 or BSA (control) as prey protein. Gel notations: (M), msGFP monomer; (D), msGFP dimer; (T), msGFP trimer; [**X**], crosslinked band representing specific docking interaction. C, densitometry analysis of specific [**X**] band fluorescence relative to total lane fluorescence normalized to the highest JNK1 concentration to create a standard curve. N=3 separate crosslinking experiments.

**Figure 3:**
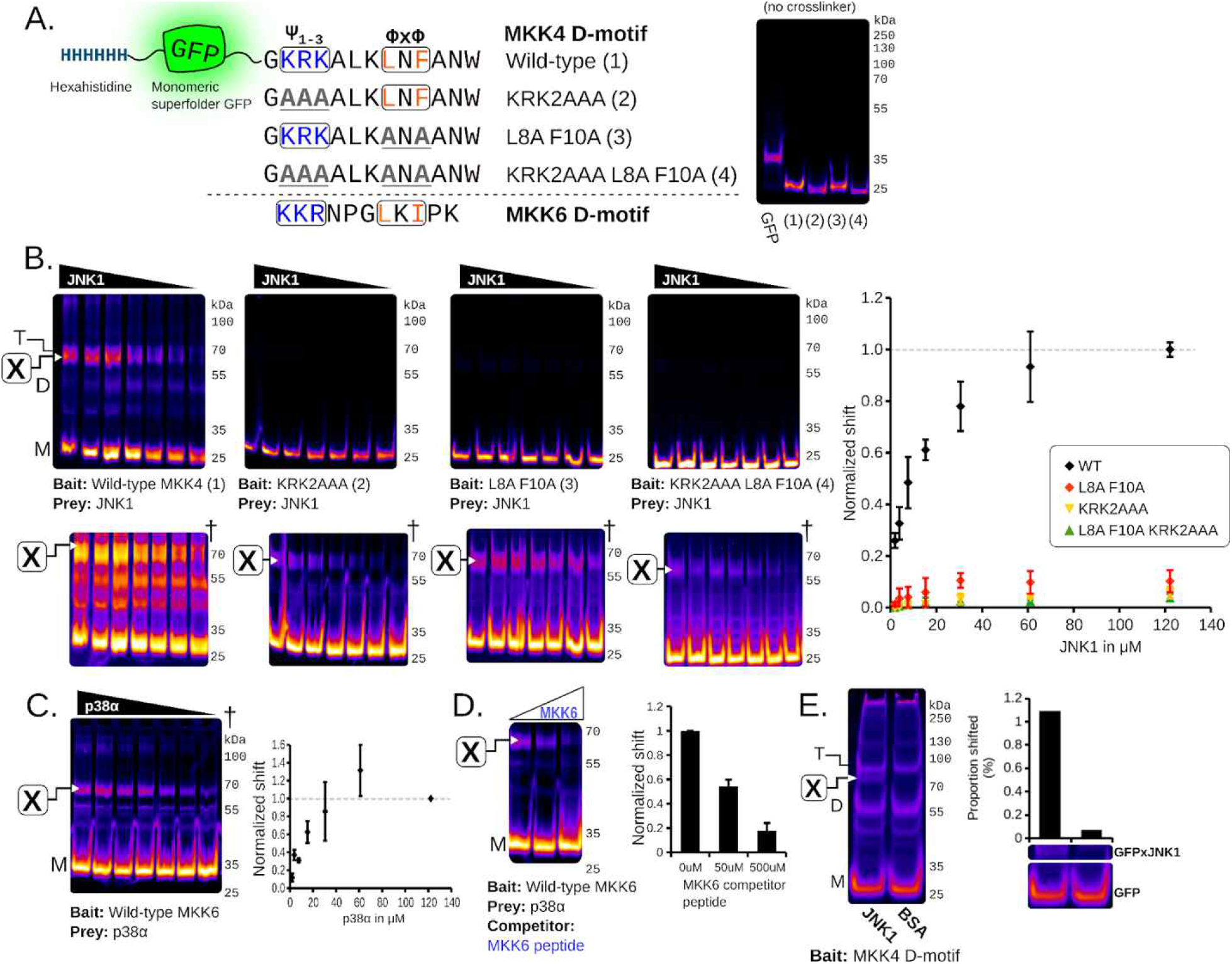
ECSA is expandable across multiple mutants and different D-motifs. A, variants of the MKK4 D-motif used to verify biochemical specificity of the assay. MKK4 peptides are labeled as (1-4) for clarity. A fluorescence gel of GFP alone along with MKK4 with crosslinker omitted is shown on the right to show differences in migration speed between the constructs. B, ECSA of wild-type and mutant MKK4 D-motifs (WT, n=3; mutants, n=2). Quantified shift is overlaid as a graph to the right and normalized to the values for the WT MKK4 D-motif. C, ECSA of the MKK6 D-motif incubated with p38α (n=2). Shift is normalized to the maximum p38α concentration to show binding curve. D, The MKK6-p38α crosslink with increasing concentrations of MKK6 D-motif synthetic peptide as a competitor, marked in blue (n=3). E, ECSA using ∼800ng of JNK1 as a prep protein showing sensitivity. For all figures, the contrast adjusted (histogram equalized) image is shown inset marked † to show the quantified bands. Binding curves of the quantified shifted band [**X**] over the total are shown at the bottom.

## Results

### Design and purification of msGFP-peptide fusions

We developed ECSA to assess interaction of peptide motifs with folded protein domains using peptide fusion to hexahistidine tagged monomeric superfolder GFP (msGFP). We opted for C-terminal msGFP fusion because some SLIMs require a C-terminal carboxyl group for interaction with cognate folded domains (7), including protein trafficking signals that are mutated in human diseases (8). This variant of GFP is highly resistant to denaturation, tolerates fusion with disordered protein sequences, and refolds rapidly after denaturation by reagents such as guanidine hydrochloride (9). The hexahistidine tag coupled with the msGFP variant allows purification of bait fusion proteins under denaturing conditions, which is helpful for peptides that are prone to aggregation or bind nucleic acids due to high positive charge.

We used the well characterized interaction of the MAPK JNK with D-motif docking peptides from MKK4 and MKK6NFAT4 to validate the ECSA approach, producing his6-tagged msGFP fused D-motif peptides (Figure 1A). To simulate conditions of semi-high throughput experiments we expressed and purified all msGFP-D motif fusion constructs in parallel from 10mL cultures (Figure 1B; see methods section); Coomassie-stained SDS-PAGE analysis of representative purity is shown in Figure 1C.

### ECSA confirms MKK4-JNK1 domain-motif interaction

We first performed experiments with msGFP C-terminally fused with the MKK4 D-motif, which like other “classical” D-motifs contains a basic region at its N-terminus and a hydrophobic (ΦxΦ) pair of residues at its C-terminus (Figure 2A). We used ECSA to evaluate this msGFP-MKK4 fusion protein with titrated JNK1, which is known to interact directly with the MKK4 D-motif. We used msGFP as a negative bait protein, and BSA as a negative control prey protein. This treatment produced a shifted band at the expected molecular weight of ∼80kDa, the combined mass of the msGFP-MKK4 fusion bait construct and the JNK1 prey (Figure 2B, left panel). Crucially, intensity of this band correlated directly with increasing concentration of added JNK1. We quantified the percent shifted in each lane using the ∼70kDa marker as a percent of the total fluorescence in that lane to determine a rough binding curve (Figure 2C). Since the maximum amount of covalently linked msGFP-protein complex is dependent on a variety of factors such as the presence of proximal lysines as well as binding affinity, we normalized the curve to the maximum bait protein concentration–in this case, JNK1. Similar amounts of BSA as well as GFP alone did not produce a band shifted species with a binding curve (Figure 2B, center and right panels).

Notably, we observed a fluorescent band at ∼55kDa in all experiments following crosslinking that was independent of prey protein presence or concentration. This occurred most dramatically in the msGFP-only experiment (Figure 2B, right panel). This is consistent with crosslinked dimers of the ∼28kDa constructs produced by oligomerization of the msGFP domains, which is present in the assay at relatively high concentration. Evidence for dimerization of msGFP has thus far only been detected by the ECSA approach.

### Analysis of ECSA specificity using mutagenesis of MKK4 D-motif regions

To assess the ECSA method’s specificity and utility for fine structure analysis of interactions between peptide motifs and folded domains, we determined if JNK1-msGFP-MKK4 D-motif crosslinking is significantly reduced by substituting amino acids in the MKK4 D-motif known to be crucial for JNK1 interaction. Specifically, we mutated MKK4 D-motif basic and hydrophobic sequences, constructing msGFP fusion proteins bearing MKK4 D-motif mutants as shown by Figure 3A (basic region, KRK2AAA; hydrophobic region, L8A F10A; both regions, KRK2AAA L8A F10A). As expected, ECSA assays with JNK1 and these mutant constructs exhibited markedly reduced production of the ∼80 kDa band-shifted species compared to the wild type, indicating a substantial loss of binding between the mutant MKK4 D-motifs and JNK1 (Figure 3B).

With enhanced contrast a faint band corresponding to the JNK1-dependent ∼80 kDa band was still observable, suggesting that binding was not entirely abolished (Figure 3B; marked †). Quantification of the abundance of this ∼80 kDa band-shifted species revealed apparent binding curves for JNK1 interaction with msGFP-MKK4 D-motif mutants KRK2AAA and L8A F10A that were considerably weaker than those observed with wild-type msGFP-MKK4. Notably, these two mutant constructs each displayed more significant JNK1 crosslinking than the KRK2AAA L8A F10A mutant construct, which eliminates both conserved basic and hydrophobic portions of the MKK4 D-motif. This mutagenesis data is consistent with previous studies using similar mutants demonstrating that the basic and hydrophobic portions of the MKK4 D-motif individually exhibit some affinity for JNK1, which employed GST-fused JNK1 in pulldown assays with 35S-labeled MKK4 (10, 11).

### Analysis of ECSA using the MKK6 D-motif – p38α interaction and peptide ligand competition

The MKK6 D-motif interaction with the MAPK p38α is similar to the interaction of the MKK4 D-motif with JNK1, with a somewhat lower affinity (3.5 vs. 7.5 μ4M). To further evaluate the utility of ECSA on a variety of peptide-protein complexes, we tested the interaction of the MKK6 D-motif with the purified MAPK p38α. As shown in figure 3C, the msGFP-MKK6 D-motif fusion protein and p38α also showed concentration-dependent production of a band-shifted species (Figure 3C). Notably, the maximum amount of shifted species was considerably weaker than the MKK4-JNK1 complex despite the comparable affinities; we speculate that this could be attributable to a lower number accessible primary amines in p38α available for crosslinking in the docked complex.

To test the specificity of the shifted ECSA band observed in the MKK6-p38α interaction, we performed a competition assay using increasing concentrations of the synthetic MKK6 D-motif (Figure 3D). Indeed, addition of either 50 or 500 μM MKK6 synthetic peptide reduced the intensity of the shifted band at the molecular weight consistent with the crosslinked species. Notably, all crosslinking reactions were performed in the presence of 20 mM primary amine (L-histidine) to quench excess crosslinker, suggesting the reduction in shift is not due to the synthetic MKK6 peptide.

### ECSA detects the MKK4-JNK1 DMI with sub-microgram prey protein input

DMIs often occur with folded eukaryotic proteins that may not purify in high yield, especially from a prokaryotic expression system. We found that JNK1-MKK4 binding could be detected at a final concentration of 0.2mg/mL, even for some MKK4 D-motif mutants, suggesting that ECSA can be sensitive enough to allow use of small amounts of input bait protein (Figure 3E).

## Discussion

ECSA is a fast, sensitive, inexpensive, and semi-high-throughput method for identifying micromolar affinity DMIs using in vitro purified components. This technique does not replace more quantitative biophysical techniques such as isothermal calorimetry (ITC) or fluorescence polarization (FP), but is rather meant to precede them to screen many DMI “hits” that merit further study. ECSA should be a useful bridge between discovery-scale experiments like phage display and more painstaking biophysical and structural characterization. The approach should prove particularly useful when identifying sites of interaction between large, disordered proteins and folded domains because it allows “scanning” of the disordered protein binding partner through rapid parallel query of many short motifs.

The peptide ligand that mediates MKK4-JNK1 interaction was first analyzed through laborious GST pulldowns with 35S-labeled MKK4 constructs (10, 11), a key initial characterization of MAPK docking (12). We used D-motif – MAPK interaction to validate ECSA. A few similar in vitro techniques exist for the discovery of weak DMIs or other PPIs. These include PUP-IT and PUP-IT2, (13, 14) which employ a bacterial enzyme, PafA, to phosphorylate a glutamate on the target peptide (Pup) fused to the bait protein. This phosphoglutamate rapidly reacts with lysine side chains on the prey to form an amide crosslink. The primary application of this technique was to detect PPIs in the cellular context, it has been demonstrated to validate multiple in vitro DMIs with low to modest affinities (14). While we have not directly compared these PafA-based approaches with ECSA, we consider them as complementary strategies.

While ECSA generates qualitative binding curves for interactions with the MKK4 and MKK6 D-motifs, the method should not be interpreted as providing quantitative data. It does not precisely mirror the in vitro binding affinities of these DMIs as rigorously determined by techniques like fluorescence polarization (FP). This discrepancy may be attributed in part to the heightened complexity of solution dynamics involving folded domains (such as JNK1 and the msGFP fusion proteins), as well as factors like the reaction of the imidoester crosslinker with the protein, protein binding and dissociation kinetics, and quenching of crosslinking by water. Notably, JNK1 interaction affinity with D-motifs has been primarily measured using synthetic peptides. It is possible that use of this peptide ligand appended to msGFP’s globular folded domain in ECSA more closely resembles the natural context of the binding interaction.

The maximal ECSA signal produced by the interaction of msGFP-MKK4 D-motif fusion with JNK1 was significantly higher than that produced by the interaction of msGFP-MKK6 D-motif with p38α, despite only a modest two-fold difference in previously measured Kd values. Consequently, the maximum attainable level of crosslinking of msGFP-peptide motif fusions with distinct prey proteins varies. Such variability is anticipated in crosslinking reactions in which primary amines must be sterically accessible within the kinetic time course of the reaction.

Despite this discrepancy in maximal signal intensity, both JNK1 and p38α D-motif interactions produced clear qualitative binding curves upon titration of the prey protein. Therefore, we recommend evaluating the concentration-dependent production of the msGFP fusion protein band-shift species by prey titration.

### Technical notes and considerations

ECSA uses an imidoester crosslinker, such as DTBP. Crosslinkers with similar linker lengths, such as dimethyl suberimidate (DMS), work as well. The approach does not work well with traditional NHS crosslinkers such as BS3, which convert the positive ε-amine of lysine into an uncharged and hydrophobic amide. In practice, we found that using NHS crosslinkers causes severe smearing on the gel, possibly due to denaturation and aggregation of the bait-prey protein complex aggravated by elimination of lysine side chain positive charge.

Purification conditions of msGFP fusion proteins bear consideration. We used a lysis buffer with 2M sodium chloride to minimize coelution of nucleic acids with basic D-motif peptides fused to msGFP. In separate experiments we found that msGFP fusion constructs containing uncharged or acidic peptides can be effectively purified using denaturing conditions (8M urea or 6M guanidinium hydrochloride), but this does not eliminate nucleic acid contamination of msGFP fusions with basic peptides such as the MKK4 D-motif (data not shown). We used L-histidine to elute msGFP-motif fusion proteins from nickel resin, which is milder than imidazole or low pH, without buffer exchange for eluted msGFP-peptide fusion proteins. We recommend alternate elution conditions or buffer exchange if prey proteins or DMIs of interest are suspected to be sensitive to L-histidine.

